# Performance characterization of PCR-free whole genome sequencing for clinical diagnosis

**DOI:** 10.1101/2020.06.19.160739

**Authors:** Ningzhi Zhang, Meizhen Zhou, Fanwei Zeng, Xiaodan Wang, Fengxia Liu, Zhihong Qiao, Chunna Fan, Yaoshen Wang, Zhonghai Fang, Wentao Dai, Jiale Xiang, Jun Sun, Zhiyu Peng, Lijie Song, Yan Sun

## Abstract

**Purpose:** To evaluate the performance of PCR-free whole genome sequencing (WGS) for clinical diagnosis, and thereby revealing how experimental parameters affect variant detection.

**Methods:** All the 5 NA12878 samples were sequenced using MGISEQ-2000. NA12878 samples underwent WGS with differing DNA input and library preparation protocol (PCR-based versus PCR-free protocols for library preparation). The DP (depth of coverage) and GQ (genotype quality) of each sample were compared. We developed a systematic WGS pipeline for the analysis of down-sampling samples of the 5 NA12878 samples. The performance of each sample was measured for sensitivity, coverage of depth and breadth of coverage of disease-related genes and CNVs.

**Results:** In general, NA12878-2 (PCR-free WGS) showed better DP and GQ distribution than NA12878-1 (PCR-based WGS). With a mean depth of ~40X, the sensitivity of homozyous and heterozygous SNPs of NA12878-2 showed higher sensitivity (>99.77% and > 99.82%) than NA12878-1, and positive predictive value (PPV) exceeded 99.98% and 99.07%. The sensitivity and PPV of homozygous and heterozygous indels for NA12878-2 (PCR-free WGS) showed great improvement than NA128878-1. The breadths of coverage for disease-related genes and CNVs are slightly better for samples with PCR-free library preparation protocol than the sample with PCR-based library preparation protocol. DNA input also influences the performance of variant detection in samples with PCR-free WGS.

**Conclusion:** Different experimental parameters may affect variant detection for clinical WGS. Clinical scientists should know the range of sensitivity of variants for different methods of WGS, which would be useful when interpreting and delivering clinical reports.

## Introduction

Massively parallel sequencing (MPS) technology is more and more widely used in genomic research and real clinical setting, which has revolutionized clinical genetic diagnosis. Recently, whole genome sequencing has been gradually implemented in the diagnosis of rare and undiagnosed clinical cases^1–3^, making it possible to be a routine in clinical care. Focusing on whole genome scale, WGS can not only be used to detect single-nucleotide variants (SNVs) and small insertions/deletions (indels), but it can also be used to identify structure variations (SVs)^4–6^. What is more, WGS can reduce the cost derived by the need of other tests^7^, and provide higher diagnostic yields than targeted panels^8^.

The process of WGS mainly includes three steps: template preparation (isolation of nucleic acid), library preparation (end repairing, adapter addition, optional PCR amplification) and sequencing (sequencing preparation, instrument operation). The results of clinical WGS may be influenced by factors related to the three steps, such as quality of genomic DNA^9, 10^, methods used for library preparation^11–13^ and differing sequencing platforms^12, 14^. After sequencing, a bioinformatics pipeline (sequencing data quality control, alignment, variant calling and interpretation) will then be implemented. The comparability of WGS can be improved by implementing a standardized bioinformatics analysis pipeline. The first three steps (template preparation, library construction and sequencing) could largely influence the quality of WGS data. As for the bioinformatics analysis of WGS, there is already a well-accepted pipeline for the analysis of SNV and indel for WGS data with short read, including alignment with BWA^15^, and variant calling with GATK^16^. There are also well-established algorithms for SV detection from short read sequencing data. Great tools and algorithms may improve the sensitivity for variant detection, however, without high quality sequencing data, it is hard to generate good results. As long as WGS data is generated, sometimes there’s little things we can do to improve the quality of WGS data. So, the performance of different experimental methods for clinical WGS and how these experimental parameters affect variant detection become interesting research topics which need to be investigated.

As for template preparation, library construction and sequencing, various methods have been provided by different sequencing platforms. Broadly, library preparation of WGS can be classified into two groups, PCR-based library preparation protocol versus PCR-free library preparation protocol. Each method has both common and specific variables related to the required DNA input, read length, and cost-effectiveness. These variables could influence the overall quality of WGS data, thus impact the sensitivity of variant detection. Specifically, in addition to evaluating the sensitivity of and breadth of coverage of WGS, we investigated the effects of library preparation method (PCR-based versus PCR-free protocols) and DNA input using MGISEQ-2000 platform in this study. In the present study, we systematically compared 5 WGS data generated from NA12878 samples. We compared the sensitivity of WGS using samples by differing library preparation protocols (PCR-based versus PCR-free protocols) and DNA inputs (1 μg, 500 ng, 300 ng and 200 ng). We also compared the yield and quality of sequencing data, depth of coverage, genotype quality, sensitivity for variant detection and breadth of coverage for each sample. The performance of each method was systematically analyzed and compared, thereby revealing how these experimental parameters affect variant detection. Generally, samples using PCR-free library preparation protocol and DNA input of 1 μg showed the highest performance in depth of coverage (DP) and genotype quality (GQ) distribution, variant detection, and breadth of coverage for disease-related genes and CNVs.

## Materials and Methods

### Samples and overall study design

To investigate the performance of PCR-free WGS for clinical diagnosis, GIAB NA12878 was collected and sequenced 5 times with differing library preparation protocols and total DNA inputs (Table 1). All the samples were sequencing on MGISEQ-2000 platform. DNA samples of NA12878 were procured from Coriell (Camden, NJ).

**Table 1.** Sample information

The overall study design is shown in Figure 1. First, to compare the performance of WGS with PCR-based and PCR-free protocols, analysis of the sensitivity of high-confidence SNPs/indels, breadth of coverage and depth of coverage were performed using PCR-based (NA12878-1) and PCR-free (NA12878-2) down-sampling samples (~40X) of NA12878. Then, using down-sampling samples of NA12878, we also compared the performance of samples (PCR-free WGS) with differing DNA input (1μg, 500ng, 300ng and 200ng) (Figure 1). This study and all the protocols were approved by the ethics committee of BGI (NO. BGI-IRB19019).

**Figure 1.**
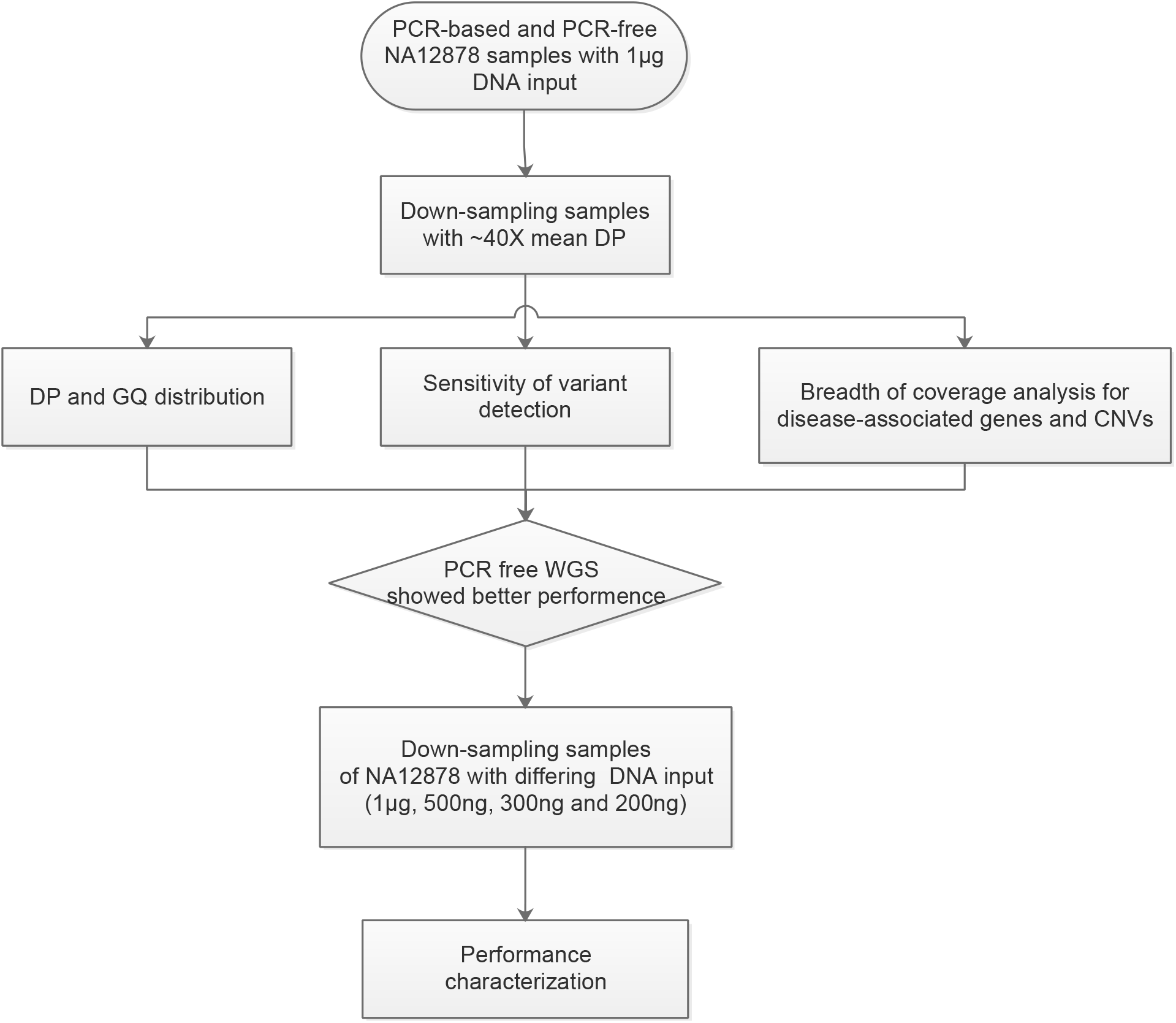
Study design

### Library preparation, genome sequencing and bioinformatics analysis

In this study, 1 μg of DNA input were used for library preparation using either PCR-based protocol (MGIEasy FS DNA Library Prep Set, containing PCR-amplification steps) or PCR-free protocol (MGIEasy FS PCR-Free DNA Library Prep Set, omitting the PCR steps). Libraries with various amount of DNA input (1μg, 500ng, 300ng and 200ng) were constructed using PCR-free library preparation protocol (Table 1). After quantification by BMG Labtech FLUOstar Omega and Agilent 2100 Bioanalyzer, all the libraries were then sequenced on the MGISEQ-2000 platform.

A standard bioinformatics analysis pipeline was implemented for all the samples. In short, after sequencing, fastq data was filtered to generate clean reads. The clean reads of each sample were then aligned to hg19 (the human reference genome) by Burrows-Wheeler Aligner (BWA)^15^. To remove duplicate reads, MarkDuplicates was then used for analysis^16^. GATK package was then used to perform realignment around indels and quality scores re-calibration, and to generate VCF files for each sample for further analysis. Depth and coverage analysis were performed by BEDTools^17^ and bamdst (https://github.com/shiquan/bamdst).

### Sensitivity and PPV of variant detection

To evaluate the performance of different experimental methods in identifying true genotypes, NA12878 high-confidence calls (v3.3.2) were recognized as true-positive calls for evaluation. We further restricted the high-confidence calls to the high confidence region to calculate the sensitivity and PPV of different experimental methods. The percentage of high-confidence calls detected by our method in all the high-confidence calls in NA12878 was considered as the sensitivity for variation detection. The percentage of high-confidence calls detected by our method in all the variants detected by our method was considered as the PPV for variation detection. To filter out erroneous variant calls, genotype quality and depth of coverage were used.

### Breadth of coverage for disease-related genes and CNVs

Genes in a single gene set may be incomplete to investigate the breadth of coverage for disease-related genes. In order to include all putative disease-related genes for evaluation, we generalize a new gene list (8,394 genes) using the following 5 databases: ClinVar (accessed on 19 February 2019), geneticHomeReference (accessed on 2 July 2019), HGMD (professional March 2018), OMIM (accessed on 4 April 2018) and Orphanet (accessed on 2 July 2019). NCBI annotation release 104 was used for the annotation of all the gene regions. Transcripts used in the HGMD database got priority for annotation. For genes without a definite transcript in the HGMD database, a combination of the regions of all transcripts was used for annotation. Coverage analysis of each sample for the 8,394 genes were performed for evaluation.

For the analysis of the coverage of CNVs, we performed coverage analysis of the NA12878 samples using CNVs from DECIPHER database (version GRCh37_v9.29).

## Results

### Overall performance of the 5 samples

The libraries of all the 5 NA12878 samples were loaded and sequenced on two lanes of MGISEQ-2000. On average, there were 256.80 Gb clean data generated per sample (two lanes). In this study, an average sequencing depth of 84.83-fold was achieved for each sample (Table 1).

In order to compare various experimental parameters (library preparation protocols and DNA inputs) at constant read depth, clean reads of each sample were randomly down-sampled from each sample using seqtk (https://github.com/lh3/seqtk). Finally, each sample was down-sampled to a sequencing depth of ~40X for further analysis.

As a result, NA12878 samples using PCR-based library preparation protocol (NA12878-1) and PCR-free library preparation protocol (NA12878-2) showed similar mean percentage of more than 98.88% and 98.62% for regions with >= 10X coverage. Samples using PCR-free WGS (NA12878-2, 3, 4 and 5) all showed a mean duplication rate of 2.5%, which was slightly lower than NA12878-1 (duplication rate of 3%).

### Distribution of DP and GQ in PCR-based and PCR-free samples of NA12878

The DP and GQ parameters are widely used for assessment of variation quality in MPS technology. In this study, we investigated the distribution of the two main quality parameters (DP and GQ) for variation detection in PCR-based (NA12878-1) and PCR-free (NA12878-2) samples of NA12878 at 1 μg DNA input. Here, down-sampling samples of NA12878 (mean DP of ~40X) was used for comparison. In general, the quality of sample with PCR-free library construction method (NA12878-2) is better than sample with PCR-based library construction method (NA12878-1) (Figure 2).

**Figure 2.**
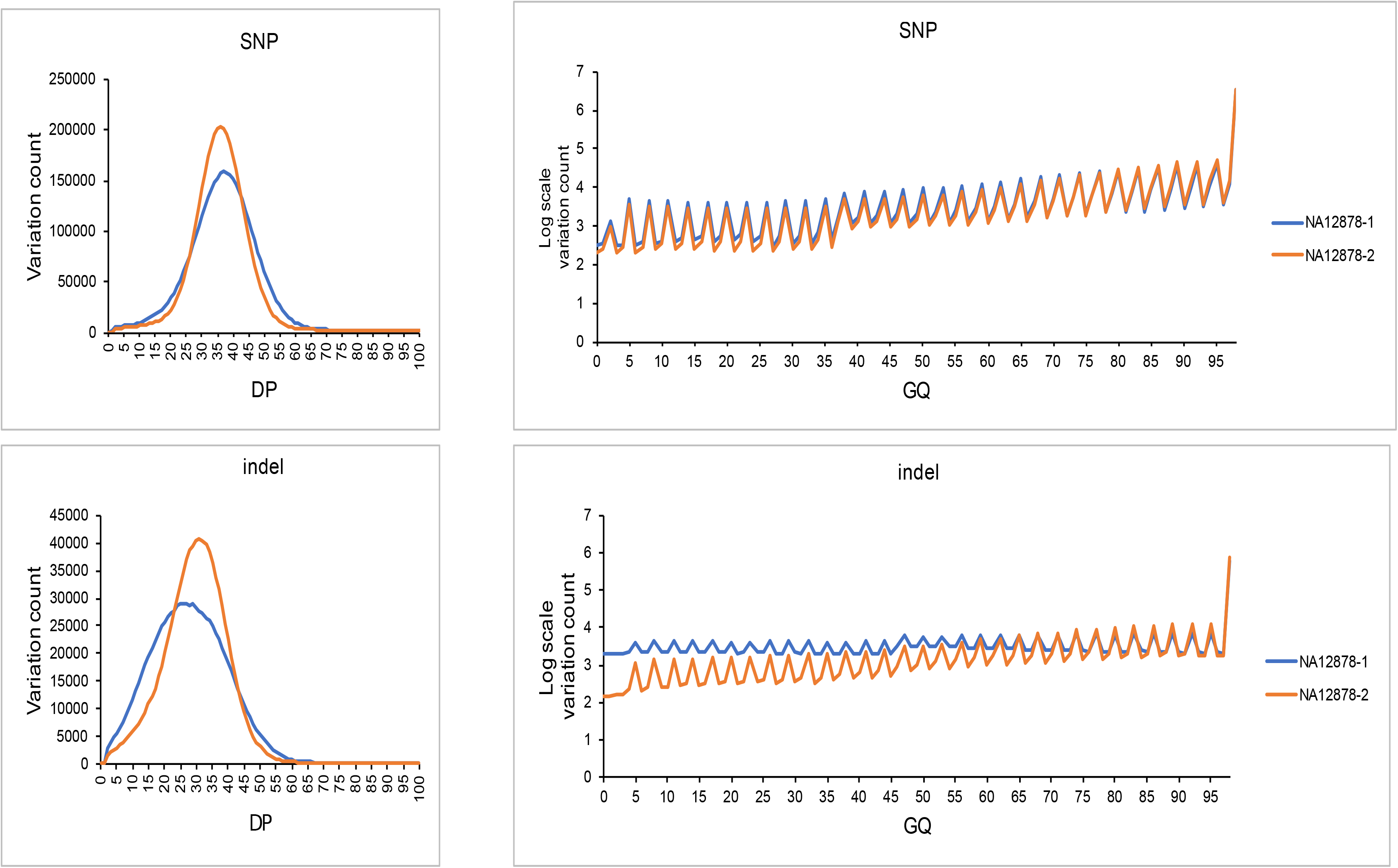
Distribution of DP and GQ in PCR-based and PCR-free samples of NA12878

The distribution of DP was normal-like for the two samples (Figure 2). The distribution of DP for all the variants showed more uniform quality for sample with PCR-free library preparation protocol (NA12878-2) than that for sample with PCR-based protocol (NA12878-1), especially for indel detection (Figure 2). The proportion of variants for NA12878-2 (93.34%) with a DP of >20X was higher than NA12878-1 (89.19%), indicating a better DP distribution for PCR-free library preparation method of WGS.

The vast majority of variants called for all the samples had a GQ close to 100 (Figure 2). Variants detected by sample with PCR-free WGS (NA12878-2) showed higher quality than those detected by PCR-based WGS (NA12878-1). The proportion of variants called for NA12878-1 (17.42%) with a GQ of <20 was 1.67% more than that for NA12878-2 (15.75%). For indel detection, more proportion of variants were detected when the GQ is less than 65 in NA12878-1 (Figure 2). These results showed that the variation quality of PCR-free WGS is better than PCR-based WGS.

### Sensitivity and PPV of variant detection in PCR-based and PCR-free samples of NA12878

In order to investigate the impact of different library preparation methods in identifying true genotypes, NA12878 high-confidence calls (v3.3.2) were recognized as true-positive calls for evaluation. To calculate the sensitivity of each sample, variants located in the high confidence region (v3.3.2) were further recognized as “gold standard” calls. To filter out erroneous variant calls, GQ (>= 20) and DP (>= 10) were used.

NA12878-2 (PCR-free WGS) showed higher sensitivity and PPV for both SNP and indel detection (Figure 3). For homozygous and heterozygous SNPs detection, the sensitivity of NA12878-2 is slightly better than NA12878-1. PCR-free WGS showed great improvement for homozygous and heterozygous indels detection (Figure 2). The sensitivity for homozygous and heterozygous indels detection in NA12878-2 is > 99.22% and 91.28% respectively, while sensitivity of NA12878-1 for homozygous and heterozygous indels detection is only 88.05% and 88.76%. For SNP and indel detection, the PPV (high confidence region) of NA12878-2 (PCR-free WGS) is also better than NA12878-1 (PCR-based WGS) (Figure 3). Heterozygous indels of NA12878-1 (PCR-based WGS) showed the lowest PPV of 82.87%. In general, the sensitivity and PPV of variant detection for samples with PCR-free library preparation protocol (NA12878-2) is better than samples with PCR-based library preparation method (NA12878-1) (Figure 3).

**Figure 3.**
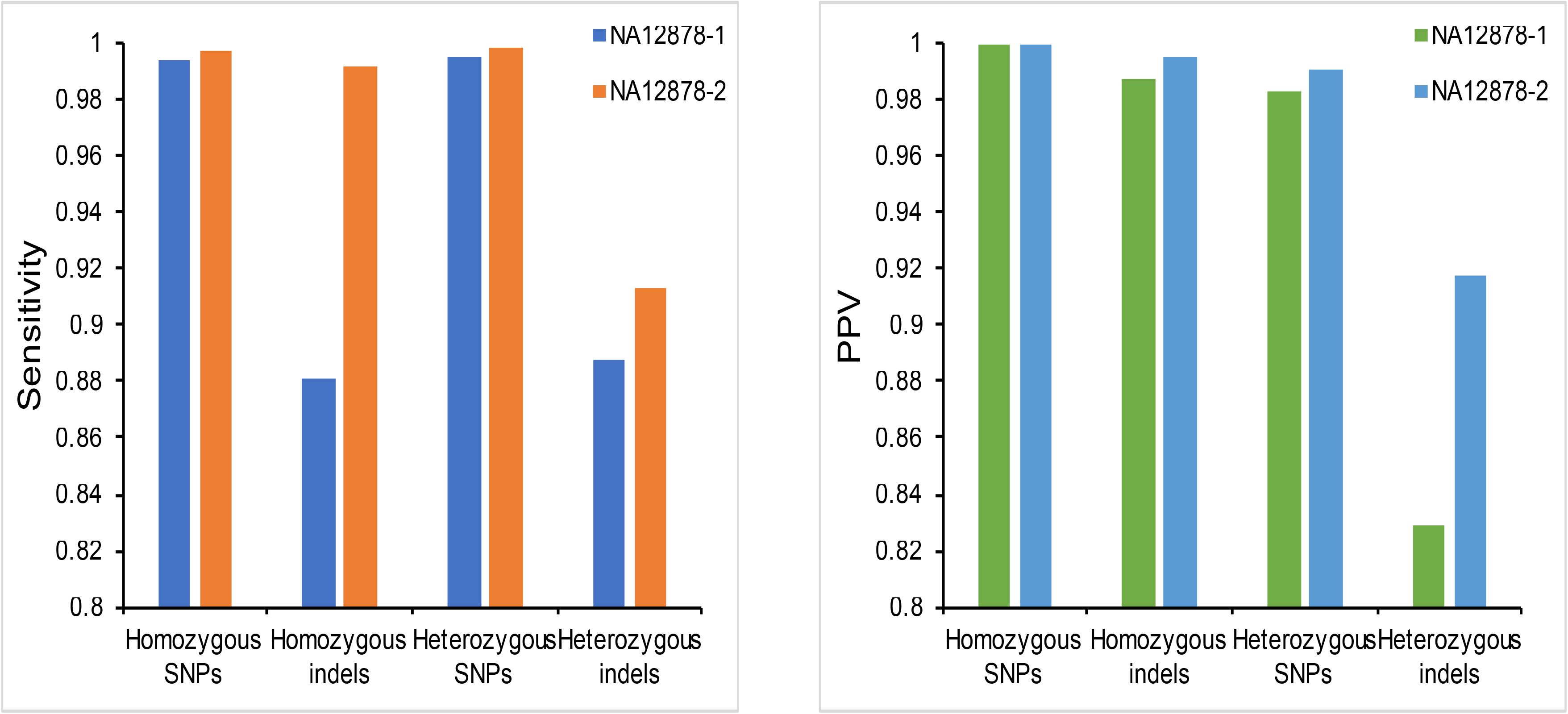
Sensitivity and PPV of variant detection in PCR-based and PCR-free samples of NA12878

### Depth and breadth of coverage for disease-related genes and CNVs in PCR-based and PCR-free samples of NA12878

In this part, the breadth of coverage for the 8,394 genes was first evaluated in PCR-based (NA12878-1) and PCR-free (NA12878-2) samples of NA12878 at 1 μg DNA input. The 8,394 disease-related genes were compiled using 5 databases (ClinVar, geneticHomeReference, HGMD, OMIM and Orphanet). The percent of targeted bases covered at > 10X depth has been reported to be related to the sensitivity for heterozygous SNV detection in WES^18^. Here, we calculated the percent of bases covered at > 10X depth for exons of the all the 8,394 genes. As a result, none of the samples of NA12878 covered 100% of the coding exons. The results of samples with PCR-free library preparation protocol method (NA12878-2) is slightly better than samples with PCR-based library preparation method (NA12878-1). For NA12878-2, the percent of bases covered at > 10X depth for the 8,394 putative disease-related genes was > 99.84%, while 99.44% of the exon regions was covered in NA12878-1.

We also compared the breadth of coverage performance of the 2 samples for ACMG 59 genes^19^. The proportion of the ACMG59 genes covered 100% (more than 10X) was 98.30% and 93.22% for NA12878-2 and NA12878-1 respectively. Sites of all genes that are covered more than 10X was 99.97% and 99.78% for sample with PCR-free library preparation method (NA12878-2) and sample with PCR-based library preparation method (NA12878-1). The breadths of coverage are slightly better for sample with PCR-free protocols (Table S1). We also examined finished genes of the ACMG 59 gene set at >= 20X coverage that could provide 99% sensitivity for heterozygous SNVs^18^. A percentage of 79.66% and 57.63% genes were covered 100% for sample with PCR-free library preparation method (NA12878-2) and sample with PCR-based library preparation method (NA12878-1) respectively.

In this study, the breadth of coverage of CNVs in the DECIPHER database was also investigated for the 2 NA12878 samples. Most CNVs in the DECIPHER database can be well covered (> 95%) at more than 10X depth for the 2 samples (Table S2). The breadths of coverage are slightly better for sample with PCR-free protocol for CNVs in the DECIPHER database (Table S2). A percentage of 92.42% CNVs showed better coverage for NA12878-2 (PCR-free WGS) than NA12878-1 (PCR-based WGS).

### Impact of DNA input in PCR-free samples of NA12878

After comparison of the performance of WGS with PCR-based and PCR-free protocols, we also investigated the impact of DNA input (1μg, 500ng, 300ng and 200ng) on variant detection in PCR-free samples of NA12878 (NA12878-2, 3, 4 and 5). We compared the DP and GQ, sensitivity for SNV/indels detection, breadth of coverage of disease-related genes and CNVs in the 4 PCR-free samples of NA12878.

First, we investigated the GQ and DP distribution in PCR-free samples of NA12878 (NA12878-2, 3, 4 and 5). In general, the distribution of DP was normal-like for all the samples. The proportion of variants with >=10X depth increased with increasing DNA input (Figure 4). NA12878-2 (1 μg DNA input) showed the highest proportion of 98.62% with >=10X depth (Figure 4). The GQ of vast majority of variants called by PCR-free samples of NA12878 (NA12878-2, 3, 4 and 5) is ~100. The proportion fluctuated along with the GQ scores. For both SNP and indel detection, the proportion of variants with >=20 GQ also increased with increasing DNA input (Figure 4). NA12878-2 (1 μg DNA input) showed the highest proportion of 99.37% with >=20 GQ (Figure 4). These results showed that the performance of samples with higher DNA input is better than samples with lower DNA input for PCR-free WGS. DNA input may influence the variant quality in samples with PCR-free WGS.

**Figure 4.**
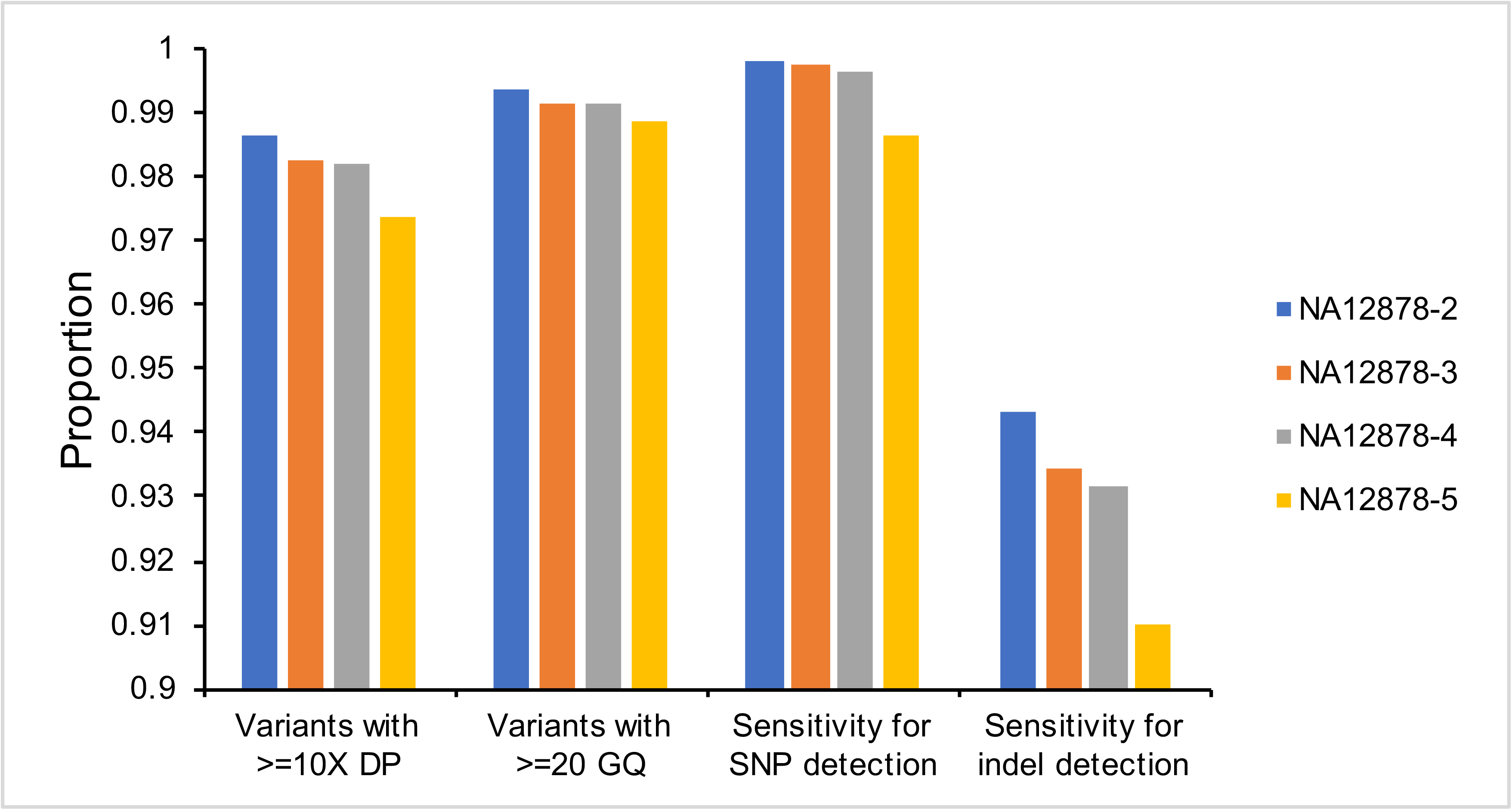
Impact of DNA input on variant detection in PCR-free samples of NA12878

DNA input may also influence the detection sensitivity of NA12878 samples. In order to investigate the impact of DNA input in identifying true genotypes, NA12878 high-confidence calls (v3.3.2) were recognized as true-positive calls for evaluation. To calculate the sensitivity of each sample, variants located in the high confidence region (v3.3.2) were further recognized as “gold standard” calls. To filter out erroneous variant calls, GQ (>= 20) and DP (>= 10) were used. With the same library preparation protocols, the sensitivity visibly increased with increasing DNA input (Figure 4), indicating that variant detection sensitivity is positively correlated with increasing DNA input in this study (Figure 4). As a result, NA12878-2 showed the highest sensitivity for both SNP (99.80%) and indel (94.34%) detection. Heterozygous indels of sample NA12878-5 (200ng DNA input) showed the lowest sensitivity of 87.92%. DNA input may influence the detection sensitivity in samples with PCR-free WGS.

We also investigated the breadth of coverage of PCR-free samples of NA12878 (NA12878-2, 3, 4 and 5) in the 8,394 disease-related genes and CNVs from DECIPHER database. To filter out erroneous variant calls, GQ (>= 20) and DP (>= 10) were used. In general, PCR-free samples with differing DNA input showed similar coverage of both putative disease-related genes and CNVs (Table S2). NA12878-5 with the lowest DNA input of 200 ng showed more lower coverage regions (Table S1, S2) in this study.

## Discussion

For specific applications of the MPS technology, PCR-based protocols are mandatory. For example, the analysis of cell free DNA. Because of its low concentration in blood plasma, PCR-based protocols are needed to provide sufficient amount of DNA for library preparation and further analysis. In this case, PCR biases would be implemented simultaneously. As is known, PCR amplification in regular library preparation of WGS causes uneven amplification and copy errors, which would result in PCR biases^11, 20–22^. PCR biases can be especially damaging to MPS applications. WGS with PCR-free protocol can remove duplicated reads generated during library construction, which can increase read utility and confidence of variant calling^23^. Other experimental parameters, such as DNA input, may also affect variant detection. However, the influence of differing experimental methods for variant detection has not been fully evaluated, especially for clinical WGS.

In this study, we focused on the impact of two experimental parameters in the upstream step of WGS analysis (library preparation protocol and DNA input) on variant detection. We comprehensively analyzed and compared the performance of each method using 5 NA12878 samples. After down-sampling to a sequencing depth of ~40X, the performance of GQ and DP for different samples were evaluated first. In addition, we further assessed the variation detection sensitivity with high-confidence calls in the high confidence region from GIAB. The breadth of coverage of disease-related genes and CNVs was also compared. As a result, samples with PCR-free protocol showed better performance in DP and GQ distribution, SNV/indel detection, and breadth of coverage of disease-related genes and CNVs, thereby revealing how experimental parameters affect variation detection. In this study, the analysis of samples with different experimental methods provided additional insight and choice for clinical variant detection.

Generally, various sequencers share a basic MPS workflow, including preparation of template, library construction, sequencing and analysis. Various experimental parameters were provided by different platforms. As for the DNA input, the data generated by extreme low DNA input may not always pass the quality control for different platforms, and that’s the reason why we selected 200 ng as the lowest amount of DNA input. The amount of DNA used for library construction can be much lower than 200 ng, such as DNA extracted from plasma and dried-blood spot. The performance characterization of extremely low amount DNA input (less than 50 ng) is another interesting research topic. Another limitation of this study is that, we did not perform CNV detection comparison in the 5 NA12878 samples, because there’s no well-accepted “gold standard” CNV call set for benchmarking, nor “best practices” workflow for the detection of CNVs. Instead, the depth and breadth of coverage for CNVs was evaluated using the 5 NA12878 samples.

The successful applications of WGS in real clinical setting requires comprehensive assessment of experimental parameters. In this study, we systematically evaluated the performance of different methods for clinical WGS, and thereby revealing how experimental parameters affect variant detection. The results provided additional insight and choice for clinical variant detection.

## Acknowledgments

We thank all the blood donors for their invaluable contribution to this study. This work was supported by the Special Foundation for High-level Talents of Guangdong (Grant 2016TX03R171). This is a non-profit research project by government.

## Availability of data and material

The data that support the findings of this study have been deposited in the CNSA (https://db.cngb.org/cnsa/) of CNGBdb with accession code CNP0001068, which is available from the corresponding author on reasonable request.

## Author Contributions

NZZ, MZZ, FWZ, LJS and YS designed the research. YS wrote the first draft of the article. XDW, ZHQ, CNF and WTD designed and performed the experiments. FXL, YSW, ZHF, JLX, JS, ZYP, NZZ, MZZ and FWZ performed data analysis. NZZ, MZZ, FWZ, LJS and WTD contributed to drafting and revising the manuscript. All authors reviewed the manuscript.

## Reference

1 Goodwin, S., McPherson, J. D. & McCombie, W. R. Coming of age: ten years of next-generation sequencing technologies. Nat Rev Genet 17, 333–351, doi:10.1038/nrg.2016.49 (2016).

2 DePristo, M. A. et al. A framework for variation discovery and genotyping using next-generation DNA sequencing data. Nat Genet 43, 491–498, doi:10.1038/ng.806 (2011).

3 Meynert, A. M., Ansari, M., FitzPatrick, D. R. & Taylor, M. S. Variant detection sensitivity and biases in whole genome and exome sequencing. BMC Bioinformatics 15, 247, doi:10.1186/1471-2105-15-247 (2014).

4 Pang, A. W., Macdonald, J. R., Yuen, R. K., Hayes, V. M. & Scherer, S. W. Performance of high-throughput sequencing for the discovery of genetic variation across the complete size spectrum. G3 (Bethesda) 4, 63–65, doi:10.1534/g3.113.008797 (2014).

5 Fang, H. et al. Reducing INDEL calling errors in whole genome and exome sequencing data. Genome Med 6, 89, doi:10.1186/s13073-014-0089-z (2014).

6 Meienberg, J., Bruggmann, R., Oexle, K. & Matyas, G. Clinical sequencing: is WGS the better WES? Hum Genet 135, 359–362, doi:10.1007/s00439-015-1631-9 (2016).

7 Soden, S. E. et al. Effectiveness of exome and genome sequencing guided by acuity of illness for diagnosis of neurodevelopmental disorders. Sci Transl Med 6, 265ra168, doi:10.1126/scitranslmed.3010076 (2014).

8 Lionel, A. C. et al. Improved diagnostic yield compared with targeted gene sequencing panels suggests a role for whole-genome sequencing as a first-tier genetic test. Genet Med 20, 435–443, doi:10.1038/gim.2017.119 (2018).

9 Zhu, Q. et al. The impact of DNA input amount and DNA source on the performance of whole-exome sequencing in cancer epidemiology. Cancer Epidemiol Biomarkers Prev 24, 1207–1213, doi:10.1158/1055-9965.EPI-15-0205 (2015).

10 Londin, E. R. et al. Whole-exome sequencing of DNA from peripheral blood mononuclear cells (PBMC) and EBV-transformed lymphocytes from the same donor. BMC Genomics 12, 464, doi:10.1186/1471-2164-12-464 (2011).

11 Aird, D. et al. Analyzing and minimizing PCR amplification bias in Illumina sequencing libraries. Genome Biol 12, R18, doi:10.1186/gb-2011-12-2-r18 (2011).

12 Clark, M. J. et al. Performance comparison of exome DNA sequencing technologies. Nat Biotechnol 29, 908–914, doi:10.1038/nbt.1975 (2011).

13 Sulonen, A. M. et al. Comparison of solution-based exome capture methods for next generation sequencing. Genome Biol 12, R94, doi:10.1186/gb-2011-12-9-r94 (2011).

14 Meienberg, J. et al. New insights into the performance of human whole-exome capture platforms. Nucleic Acids Res 43, e76, doi:10.1093/nar/gkv216 (2015).

15 Li, H. & Durbin, R. Fast and accurate short read alignment with Burrows-Wheeler transform. Bioinformatics 25, 1754–1760, doi:10.1093/bioinformatics/btp324 (2009).

16 McKenna, A. et al. The Genome Analysis Toolkit: a MapReduce framework for analyzing next-generation DNA sequencing data. Genome Res 20, 1297–1303, doi:10.1101/gr.107524.110 (2010).

17 Quinlan, A. R. & Hall, I. M. BEDTools: a flexible suite of utilities for comparing genomic features. Bioinformatics 26, 841–842, doi:10.1093/bioinformatics/btq033 (2010).

18 Kong, S. W., Lee, I. H., Liu, X., Hirschhorn, J. N. & Mandl, K. D. Measuring coverage and accuracy of whole-exome sequencing in clinical context. Genet Med 20, 1617–1626, doi:10.1038/gim.2018.51 (2018).

19 Kalia, S. S. et al. Recommendations for reporting of secondary findings in clinical exome and genome sequencing, 2016 update (ACMG SF v2.0): a policy statement of the American College of Medical Genetics and Genomics. Genet Med 19, 249–255, doi:10.1038/gim.2016.190 (2017).

20 Kebschull, J. M. & Zador, A. M. Sources of PCR-induced distortions in high-throughput sequencing data sets. Nucleic Acids Res 43, e143, doi:10.1093/nar/gkv717 (2015).

21 Chen, Y. C., Liu, T., Yu, C. H., Chiang, T. Y. & Hwang, C. C. Effects of GC bias in next-generation-sequencing data on de novo genome assembly. PLoS One 8, e62856, doi:10.1371/journal.pone.0062856 (2013).

22 Ross, M. G. et al. Characterizing and measuring bias in sequence data. Genome Biol 14, R51, doi:10.1186/gb-2013-14-5-r51 (2013).

23 Jones, M. B. et al. Library preparation methodology can influence genomic and functional predictions in human microbiome research. Proc Natl Acad Sci U S A 112, 14024–14029, doi:10.1073/pnas.1519288112 (2015).

